# Recurrent *AIPL1* c.487C>T truncating variant in Leber Congenital Amaurosis: Support of pathogenicity and regional implications

**DOI:** 10.1101/290650

**Authors:** Mohammed O.E. Abdallah, Mahmoud E. Koko, Shima Faisal, Melanie J. Newport, Muntaser E. Ibrahim

**Affiliations:** Department of Molecular Biology, Institute of Endemic Diseases, University of Khartoum, Khartoum, Sudan; Department of Neurology and Epileptology, Hertie Institute for Clinical Brain Research, Tuebingen, Germany; Faculty of Pharmacy, University of Khartoum, Khartoum, Sudan; Wellcome Trust Brighton and Sussex Centre for Global Health Research, Brighton and Sussex Medical School, Brighton, UK

**Author notes:** Corresponding author: Prof. ME Ibrahim, Institute of Endemic Diseases, Department of Molecular Biology, University of Khartoum, 11111 Khartoum, Sudan. P.O. box 102. Equal contribution (Mohammed O.E. Abdallah, Mahmoud E. Koko, Shima Faisal).

**Keywords:** Leber Congenital Amaurosis, LCA4, AIPL1 c.487C>T, Sudan, AR autosomal recessive

## Abstract

**Background:** Leber Congenital Amaurosis (LCA) is a clinically and genetically heterogeneous inherited retinal dystrophy characterized by early onset visual impairment caused by mutations in not less than 17 genes. *AIPL1* mutations cause LCA type 4, comprising approximately 7% of LCA worldwide. The importance of establishing a genetic diagnosis lies in the promise of gene therapy demonstrated in mouse models.

**Results:** we genetically investigated a consanguineous Sudanese family with Leber Congenital Amaurosis. Eight members of the family were affected. Using whole exome sequencing in two siblings and their healthy mother, both inheritance-based and phenotype-based prioritization strategies converged to identify a truncating variant (rs62637009) in *AIPL1*, consistent with a diagnosis of LCA type 4. *AIPL1* c.487C>T is an ultra-rare cause of LCA4 that was seen previously in homozygous state in a single Palestinian family. This recurrent variant seems to have a regional importance with a likely founder effect.

**Conclusions:** This report adds evidence to the pathogenicity of *AIPL1* c.487C>T meriting its conclusive annotation as a recurrent pathogenic variant. This variant is particularly relevant to the middle-eastern and northeast African regions.

## Introduction

Leber Congenital Amaurosis (LCA) is a clinically and genetically heterogeneous inherited retinal dystrophy characterized by early onset visual impairment and nystagmus. The diagnosis of LCA is usually clinical. Pathogenic variants in 17 genes are known to cause LCA and at least one other disease locus for LCA has been reported [1]. These genes encode a variety of proteins, including those involved in developmental and physiological pathways in the retina [2]. LCA type 4 (LCA4) is caused by *AIPL1* gene located in chromosome 17p13.1. *AIPL1* mutations cause approximately 7% of LCA worldwide. It was first identified in a Pakistani family with autosomal recessive amaurosis [3].

Mutations in *AIPL1* are associated with a form of LCA that is characterized by maculopathy and a pigmentary retinopathy starting at a young age [4]. AIPL1 (aryl hydrocarbon receptor-interacting protein-like 1) which is expressed in rod and cone photoreceptors, has a critical role in cell viability and the assembly of the phototransduction protein, phosphodiesterase, in both rods and cones [5]. In mouse models of AIPL1 deficiency, rapid complete degeneration of both rods and cones is seen within 4 weeks of age [6, 7]. Establishing a genetic diagnosis for LCA is of great importance as the prospects of gene therapy seems promising, especially for LCA caused by AIPL1 as demonstrated in mouse models [8].

Multiple genetic variations in *AIPL1* has been reported to cause of LCA4. As of the end of September 2017, the ClinVar [9] database lists 10 pathogenic and likely pathogenic variants in *AIPL1* including one truncation (p.(Trp278Ter)), two insertion-deletions (p.(Glu337Alafs) and p.(Al352_Pro355del)), and six missense (p.(Ala197Pro), p.(Ile206Asn), p.(Cys239Arg), p.(Gly262Ser), p.(Arg302Leu), p.(Pro376Ser)) variants, while more than 50 variants are reported in the HGMD database [10]. Here we report the first Sudanese family with autosomal recessive Leber Congenital Amaurosis resulting from *AIPL1* c.487C>T (p.(Glu163Ter)) truncating variant (rs62637009) diagnosed using whole exome sequencing. To our knowledge, this is the first phenotype-genotype correlation of *AIPL1* with LCA4 in the east African region, and the second report of c.487C>T worldwide that was originally seen in a Palestinian family [11].

## Methods

We genetically investigated two siblings with early onset visual impairment and a family history of a similar condition using whole exome sequencing. The first patient was a female aged twenty-four years, who was blind at time of investigation. The onset of visual impairment was noted since birth with severe loss of vision in early childhood (Acuity: Perception of Light). Photo-attraction was present (staring at light) as well as retinal pigmentation. She developed keratoconus and cataract resulting in complete blindness. Her younger brother (six years old) had a similar clinical presentation (acuity: Perception of Light). The family pedigree showed another affected sibling along with five related patients with similar phenotypes. Figure (1) shows the family pedigree.

### Whole exome sequencing

Whole exome sequencing was performed on two affected probands and their mother. Paired-end sequencing was done on Illumina HiSeq platform (Illumina, CA, USA) using DNA extracted from peripheral blood samples. Agilent SureSelect enrichment kits were used (Agilent Technologies, CA, USA). Following alignment with BWA MEM [12], sorting and duplicates removal with Samtools [13], variants were called using Freebayes [14]. Variant calls were annotated using SnpEff v4.1 [15] and Ensembl VEP v86 [16]. The annotations included ExAC allele frequencies [17] and multiple pathogenicity predictions.

### Linkage-based Variant Prioritization

the consistency of the phenotype, its severity along with the consanguinity seen in the pedigree were highly suggestive of the inheritance of a rare or novel variant in a homozygous region. Gemini v0.18 [18] was used for identification of rare homozygous variants segregating in both patients. We performed homozygosity mapping using Homozygosity Mapper [19] and looked for homozygous runs shared by both patients. All regions with scores above 90% of the maximal exome wide homozygosity signal were prioritized. After the exclusion of the homozygous runs seen in the mother, we examined all the rare or novel variations in shared runs. Figure 2 Shows

**Figure 1.**
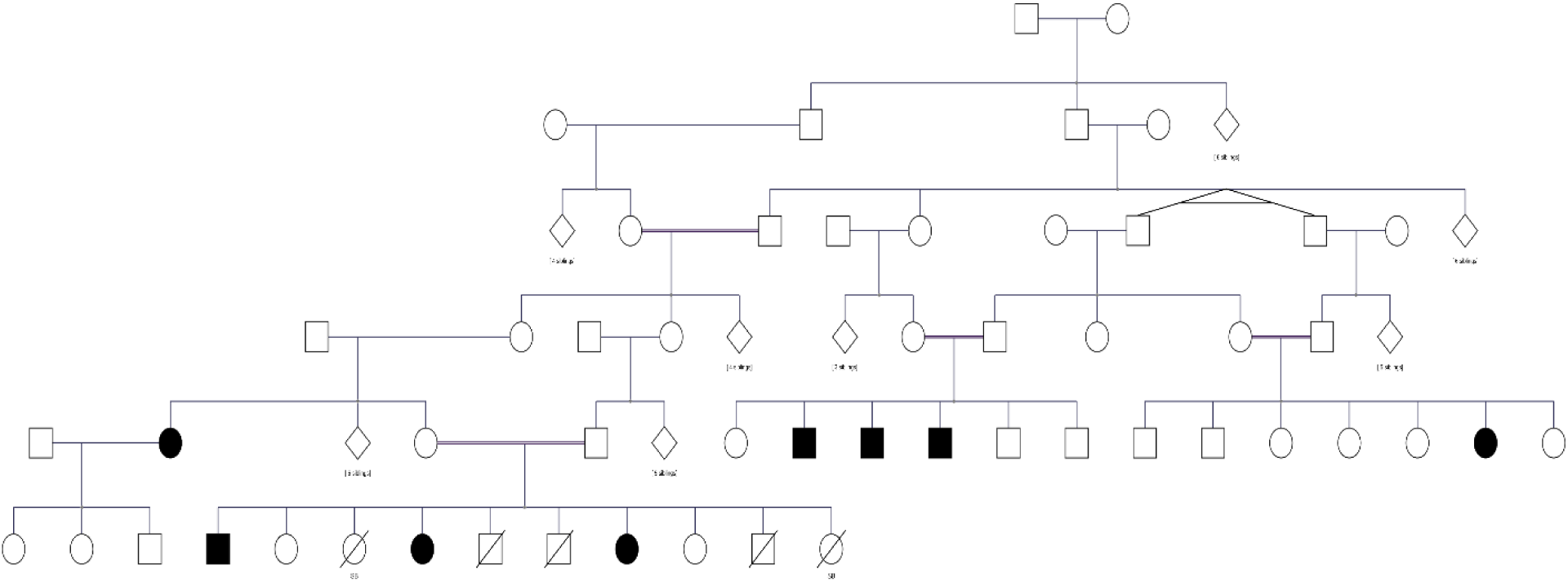
the pedigree of consanguineous multigenerational family with Leber congenital amaurosis.

**Figure 2.**
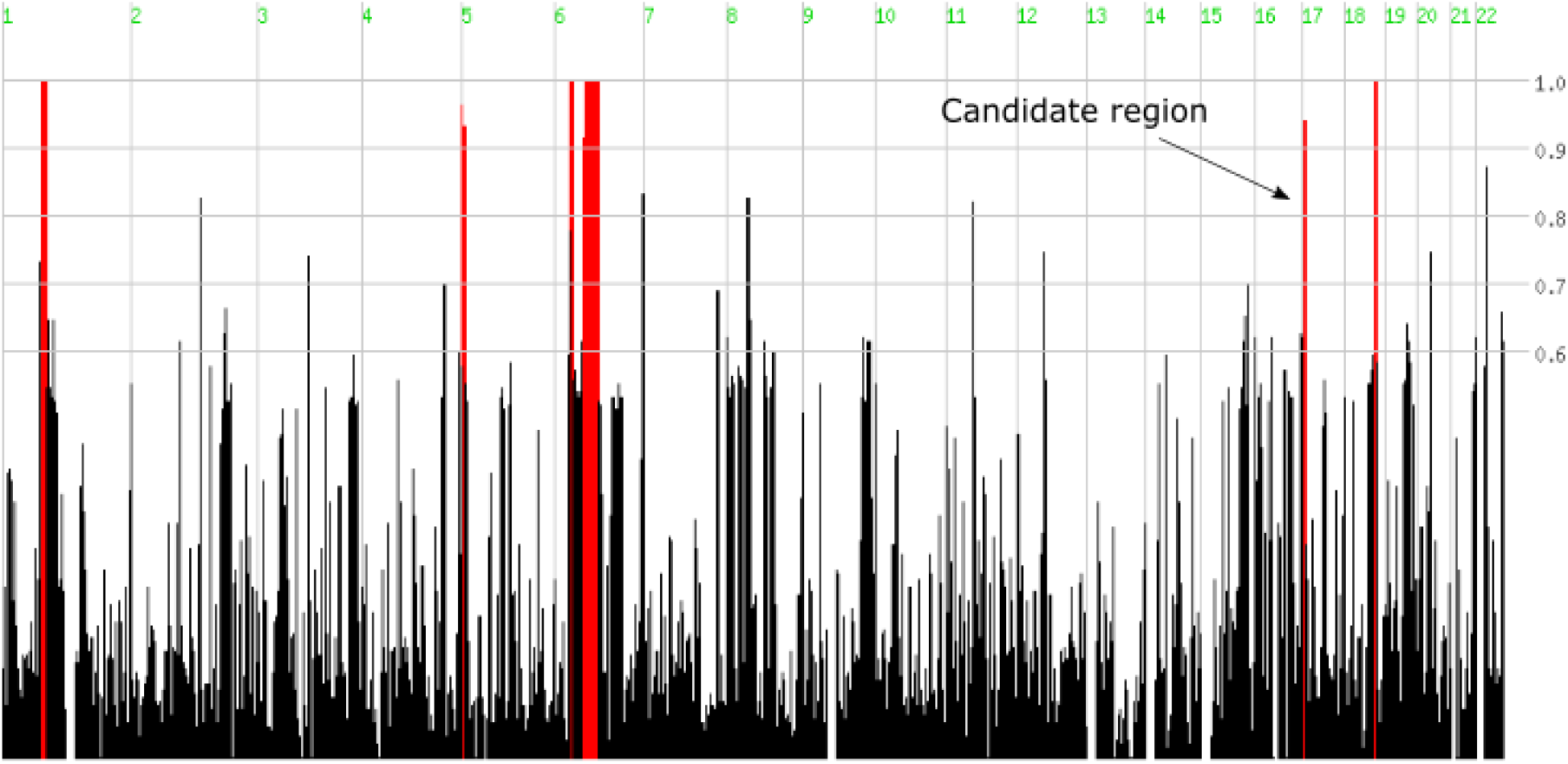
Illustrates runs of homozygosity (ROHs) identified by HomozygosityMapper.

### Phenotype-based Variant Prioritization

we obtained a list of genes linked to LCA in OMIM [20] database under the phenotypic series PS204000, which included seventeen genes. OpencgaR v1.3 (https://github.com/opencb/opencga/) was used to mine all variants overlapping this gene list from the VCF files. Scaled CADD[21] score cutoff of 20 was used for deleteriousness filtering. All variants exceeding this score were manually evaluated for their pathogenicity and relation to the phenotype. We also used Exomizer v8.0.1 [22] for automated variant prioritization (complete results and parameters for Exomizer analysis are available in supplementary materials file S1).

## Results

### Inheritance-based variant prioritization

The highest shared homozygosity signals were seen on chromosomes 5, 6 (two runs of homozygosity), 17, and 18 with a length of 0.54 Mb, 9.4 Mb, 15.3 Mb, 0.6 Mb, and 1.6 Mb, respectively. We excluded a region in chromosome 1 that was similarly homozygous in the mother. In total, only four rare variants were seen in these regions that fit a recessive inheritance pattern. The variant rs62637009 was the only pathogenic variant (stop gain). This truncating variant in *AIPL1* (NM_014336.4:c.487C>T) was in a homozygous state in the patients and heterozygous carrier state in their mother with coverage exceeding 20x in all three samples. It is predicted to introduce an early stop codon (NP_055151.3:p.(Gln163*))(Figure (3). The remaining three variants (rs759751696, rs2089318, rs144622175) were all predicted to have a non-coding modifier effect. It is worth mentioning that the length of the homozygous run did not indicate the position of the causative variant.

### Phenotype-based variant prioritization

92 variants in LCA genes were detected, including four variants that followed an AR pattern of inheritance (rs62637009, rs2297129, rs2297128, and rs2274736). Only rs62637009 had a deleterious scaled CADD score above 20 (36). Interestingly, rs62637009 was also the top hit in Exomizer results, with a score of 0.818, a phenotype score of 0.584, and a variant score of 0.950 Clinical Interpretation of rs62637009 according to ACMG/AMP 2015 guidelines [23] Using ACMG wIntervar [24] this variant was classified as pathogenic on the basis of the following evidence codes: PVS1, PS1, PM2, and PP3 (null variant (nonsense), Same amino acid change as a previously established pathogenic variant regardless of nucleotide change, Absent from controls (or at extremely low frequency if recessive) in Exome Sequencing Project, 1000 Genomes Project, or Exome Aggregation Consortium, multiple lines of computational evidence support a deleterious effect on the gene or gene product)

## Discussion and conclusions

Identification of rare disease variants is critical for the accurate diagnosis, counseling and any future gene-specific therapeutic interventions. This presents an everyday challenge for clinicians and geneticists. Traditional sequencing of a limited number of genes is a strategy that fails to cope with the genetic heterogeneity seen in many disorders. Using population-optimized microarrays is yet another proposed solution that will likely struggle to accommodate the expanding number of known and novel variants with different frequencies that probably relate to each population’s structure and evolutionary history. As well, when assessing an individual patient, the answer to the question ‘which population is representative of each patient’ might not be as clear as we expect.

Exome sequencing provides an exceptional opportunity to identify ultra-rare and novel variants in single families. However, the challenge of variant prioritization in exome sequencing studies is a big hurdle. In this study we demonstrated that employing an integrative approach combining genotype-based strategies like homozygosity mapping (in consanguineous families) and phenotype-based approaches, e.g. through Exomizer has the potential to converge on the same causative variant. While homozygosity mapping continues to prove its worth in identifying recessive disease variants, phenotype-based strategies coupled with deleteriousness score filtering are suitable for many scenarios of Mendelian and non-Mendelian inheritance.

We identified two patients who harbor a rare homozygous nonsense variant in *AIPL1* that has been previously reported only once in homozygous state [25]. The patients in the previous report by Tan *et al.* [11] were of Palestinian origin, highlighting the importance of extremely rare, population-specific disease-causing variants. This variant was not detected in many large databases including ExAC [17] and GnomAD indication a possible founder effect. The phenotype observed in our patients matched the phenotype previously described for this type of LCA with an early onset loss of vision characterized by a very low visual acuity (light perception) that further deteriorates following the development of keratoconus and cataracts as well as pigmentary retinopathy, also characteristic of AIPL1 forms of LCA [26]. Assessing this variant against the ACMG guidelines [23] clearly pinpointed its clinical significance, adding further evidence to its pathogenicity.

This variant is located distal to the FKBP-type peptidylprolyl isomerase domain and before the TPR domain (formed of Tetratricopeptide repeat-containing domain, Tetratricopeptide-like helical domain and a Tetratricopeptide repeat motif) and C-terminus Proline-rich domain. TPR domain mediates protein-protein interactions and the assembly of multiprotein complexes [27]. Variants affecting the TPR domain of AIPL1 are thought to disrupt its interaction with Hsp90 and Hsp70 [28] preventing the formation of a retina-specific chaperone heterocomplex facilitating retinal protein maturation[28]. This variant results in a short truncated protein with loss of terminal domains, and is most likely eliminated by Nonsense-mediated decay (NMD), although this requires experimental validation. However, early null variants typically cause NMD with a subsequent protein deficiency. Figure 3 shows a lollipop plot of AIPL1 gene variants

**Figure 3.**
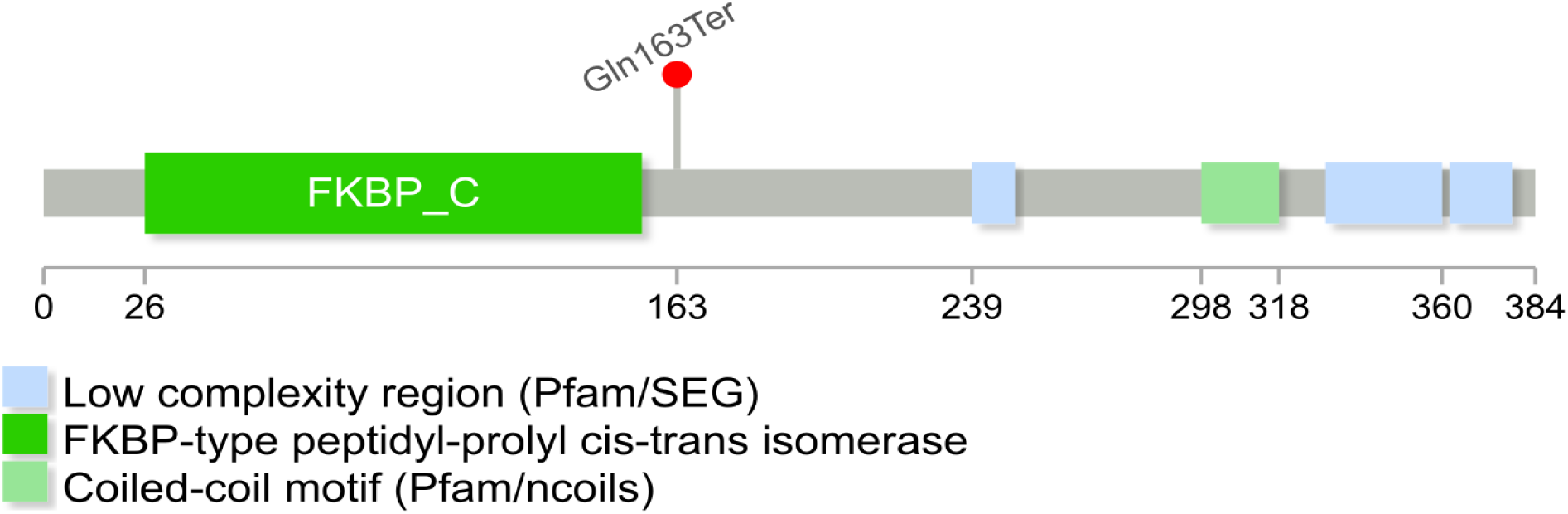
a lollipop plot that illustrates the impact of rs62637009 on the AIPL1 protein.

In summary, there is enough evidence to implicate NM_014336.4(AIPL1):c.487C>T as a pathogenic ultra-rare truncation causing Leber Congenital Amaurosis with rapid and severe progression. This variant also seems to have a regional significance with a possible founder effect.

## List of Abbreviations

Variant, NMD, LCA

## Declarations

### Ethical approval and consent for publication

was obtained from the institutional ethical committee of the Institute of Endemic Diseases, University of Khartoum. Informed consent was obtained from adult participants and the parents of participating children for the genetic analysis, the publication of this report and all accompanying materials. All investigated individuals agreed to the publication of this report including their anonymous clinical data, family pedigree and genetic results.

### Availability of data and materials

Supplementary file (I) is Exomizer results file. The raw exome datasets analyzed during the current study are not publicly available to abide to the confidentiality requested by the participants.

### Authors’ Contribution

SF, MI and MN conceived and designed the study. SF obtained the data and processed the blood samples. MI and MN facilitated the exome sequencing. MA and MK analyzed the exome data and interpreted the results. MK and MA wrote the manuscript with critical revisions from SF, MI and MN. All authors read the manuscript and approved its final version.

## Acknowledgments

Not applicable.

## Supplementary Material

File S1 Exomizer results and analysis paramters

